# Urban bats change the menu: dietary plasticity across human-modified landscapes of a tropical island

**DOI:** 10.1101/2025.05.06.652573

**Authors:** Muriel Dietrich, David A. Wilkinson, Gildas Le Minter, Magali Turpin, Maxime Galan, Nathalie Charbonnel, Guillaume Dupuy, Nausicaa Habchi-Hanriot, Bernard Reynaud, Hélène Delatte, Camille Lebarbenchon

**Author notes:** UMR ASTRE (Animal, Santé, Territoires, Risques, Ecosystèmes), CIRAD / INRAE / Université de Montpellier, 97490 Sainte-Clotilde, Reunion Island. **Correspondance** Muriel Dietrich - UMR PIMIT, Sainte-Clotilde, Reunion Island.

## Abstract

In a rapidly urbanizing world, research on dietary habits of wildlife is essential to understand how plastic behaviors may guide evolutionary trajectories of endangered animal species. Bats are very sensitive to habitat destruction and land-use change, although some species have shown the capacity for adaptation to urban life. In this study, we tested to what extent urban insectivorous bats modify their feeding strategies in a recently human-modified tropical insular ecosystem. Using a DNA metabarcoding approach on fecal samples collected in seven roosts, we analyzed the dietary niche of free-tailed bats (*Mormopterus francoismoutoui*) endemic to Reunion Island. Our results revealed a wide dietary niche, including 174 arthropod species in 12 orders, among which lepidopterans were predominant. We identified several crop pests and disease vectors, highlighting the central role of this bat species for agroecology and epidemiology concerns. Our study also highlighted potential sex- and reproduction-related dietary strategies. Moreover, we found that agriculture areas, inferred from land cover surrounding bat roosts, were associated with higher relative abundance of Lepidoptera in the diet of bats. In contrast, bats roosting in urban areas increased their consumption of Blattodea. As Reunion free-tailed bats roost and thrive in human-modified landscapes, understanding the consequences of this dietary plasticity for bat health and fitness will be necessary for urban evolutionary research and conservation actions.

## 1 INTRODUCTION

Urban and agricultural land expansions are considered to be one of the main causes of environmental changes at the global scale [1]. Today, farmlands dominate 38% of the global land surface [2] and, by 2070, 58% of the human population is projected to live in an urban area [3]. These profound human perturbations can decrease species richness and modify community composition, with potential impacts on resource availability and diet selection for predators [4]. This, in turn, can have detrimental effects on body condition, physiology, reproduction, and finally fitness of individuals [5], leading gradually to the decline of animal biodiversity. Yet, recent studies show that some species adapt their behavior and may adjust their foraging strategies to novel environments [6–8]. Such urbanization-associated dietary plasticity can have both positive and negative consequences for wildlife health and ecology [9,10]. Research on feeding behavior is therefore essential to understand whether and how plasticity may guide eco-epidemiological processes and evolutionary trajectories in human-modified landscapes [11].

Bats are among the most common mammals in urban habitats [12] but they face many threats which reduce their populations worldwide [13]. Bat feeding strategies are a very important part of their ecology [14]. Indeed, because of exceptionally high energy costs during active flapping flight, insectivorous bats may consume up to 80% of their body mass each night [15,16]. With such high food intake, insectivorous bats are thus recognized as main biological suppressors of arthropods, including pests and disease vectors [17–21]. However, insectivorous bats have to face the global decline of insect abundance and diversity, for which urbanization and agriculture are major drivers [22]. Dietary plasticity of bats to human-modified landscapes has commonly been examined at the species level and several studies have shown that bats may adapt their diet, through ecological and evolutionary processes, depending on the species-specific roost environment and energetic requirements [23–25]. For example, agriculture reduces prey diversity of the European insectivorous bat *Miniopterus schreibersii*, which feeds on moths in surrounding crops [26]. Moreover, sex- and reproduction-specific diets have been observed in bats (e.g. [27,28]). These patterns might reflect habitat selection based on different energy requirements, such as for females of *Myotis dasycneme* in the Netherlands, which predate more abundant but lighter prey when pregnant [29]. Around the world, molossid bats are the dominant species within urban bat assemblages, frequently found foraging and roosting in urban areas, and are thus considered well-adapted to human-modified landscapes [30]. However, little is known on how they cope with land-use changes and if sexes may adapt differently their dietary niche.

The Reunion free-tailed bat (*Mormopterus francoismoutoui*) is a tropical insectivorous molossid bat endemic to Reunion Island [31]. This small volcanic territory is located in the south-western Indian Ocean (Mascarene Archipelago) and is a major biodiversity hotspot [32]. However, since its first human colonization 350 years ago, natural ecosystems of Reunion Island have suffered from deforestation, agricultural expansion and urbanization [33,34]. New ecological and societal challenges arise with the recent biological invasions of crop pests and disease vectors, the respective management of agricultural production [35] and the emergence of human and livestock diseases, such as dengue and bluetongue [36,37]. Despite rapid landscape modifications in Reunion Island, numerous free-tailed bats show opportunistic behaviors and roost in urban areas, such as in buildings and bridges, suggesting some plasticity in their foraging strategies [31,38,39]. Although the Reunion free-tailed bat is widespread on the small territory of Reunion Island, its biology has only recently been documented [39,40] and its dietary niche has not been characterized so far. The Reunion free-tailed bat thus provides an original and striking model for understanding urban plasticity of bats in human-modified landscapes.

The objectives of our study were (i) to assess the realized dietary niche of free-tailed bats throughout Reunion Island, including arthropod pests and disease vectors, (ii) to test for roost, sex- and reproduction-related variations in dietary niche, and (iii) to determine to what extent agriculture and urbanization affect dietary composition. We specifically tested whether the extension of agricultural and urban areas would reduce the dietary niche of bats, as a consequence of potential insect decline.

## 2 MATERIALS AND METHODS

### 2.1 Field sampling

We collected fecal samples from *M. francoismoutoui* during summer months in 2018 (from October 29^th^ to December 7^th^) at seven roosts located all over the island (Figure 1a). We choose this time frame as it corresponds to the gestation in females and could lead to increased sex-specific foraging behaviors, due to the energetic requirements of pregnant females [28]. Each roost was sampled once and one fecal pellet was analyzed per bat individual. Bats were captured at night after dusk emergence, using harp traps installed close to the roost exit, without preventing the exit of bats. Due to the accessibility of one roost, a butterfly net was used instead by carefully approaching resting (non-flying) individuals during the day. After capture, bats were immediately hydrated with a sterile syringe and water, placed in a clean individual bag close to a warm source (hot water bottle), and processed at the capture site. Fresh feces samples were collected either from the bag or directly when handling bats using cleaned tweezers, and stored in a sterile vial in an ice cooler in the field, prior to transfer to a -80°C freezer. We also visually determined the sex and reproductive status of each individual (all bats were adults) as previously described [39]. Pregnancy was determined by slight palpation of the abdomen and the presence of enlarged nipples. Finally, each individual bat was immediately released on the capture site after being processed.

**Figure 1.**
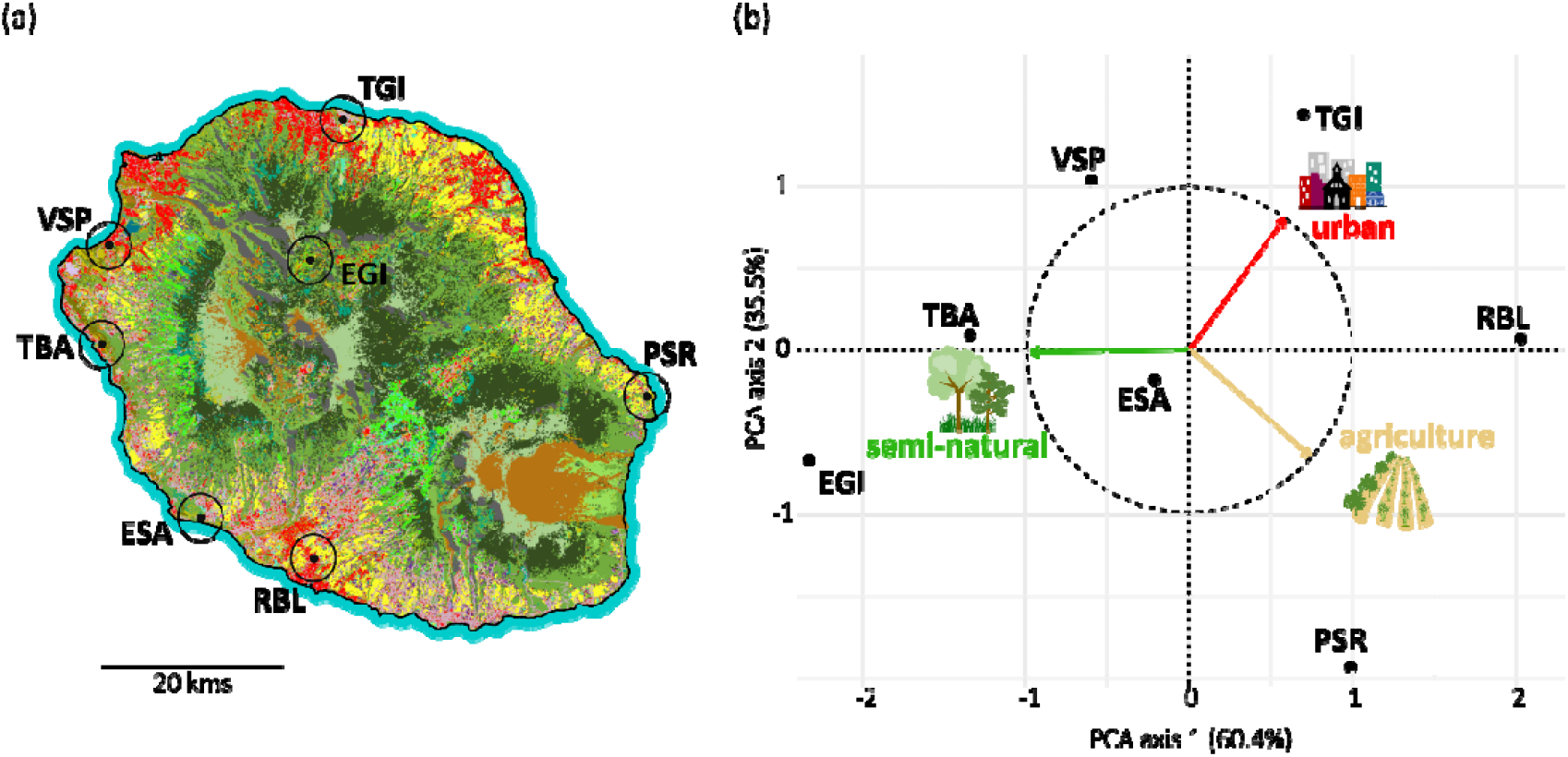
Location of bat roosts and Principal Component Analysis across different landscapes in Reunion Island. (a) Map of Reunion showing habitat types (detailed in Figure S1), with roosts represented by small black points labeled with three-letter codes. Black circles indicate the landscape buffer zone considered for each roost. (b) PCA plot illustrating the distribution of roosts based on three land-use variables (arrows).

Handling of bats was performed using personal protective equipment (FFP2 masks, nitrile gloves, as well as Tyvek suits and respirator cartridge masks inside the cave). Gloves were disinfected between each bat individual and changed regularly, and all the equipment was disinfected between roosts as well.

### 2.2 Landscape characterization

Land-use data of Reunion Island in 2021 was retrieved from the CIRAD “aware” website, which includes 28 different habitats. We quantified landscape composition around the sampled roosts, by overlaying a 5-km buffer zones over each roost location (Figure 1a), which should encompass the habitat in which bats forage around the roost, based on preliminary tracking data on the hunting distance of Reunion free-tailed bats [41]. The area of the 28 habitats was obtained for each roost and aggregated into three land-use variables: urban, semi-natural and agricultural areas (habitats included in each category are listed in Figure S1). To visualize landscape differences among roosts, a Principal Component Analysis (PCA) was applied on the three land-use variables, using the *FactoMine* R package [42]. A landscape dissimilarity matrix among samples, based on the three land-use variables, was then build using Euclidean distances and the *vegdist* function in R-4.4.2.

### 2.3 DNA extraction and PCR amplification

DNA was extracted from one fecal pellet per bat resuspended in 700 μL of MEM eagle. Then, 200ul of sample was used with the Cador Pathogen 96 Qiacube HT kit (Qiagen), and each run included a negative extraction control. We amplified a 157-bp COI minibarcode using the common ZBJ primers [43]. The PCR was performed in 22 μL reaction volume using 10 μL of Gotaq® G2 Hot Start Green Master Mix 2X (Promega), 1 μL of each primer (10μM), and 10 μL of DNA extract. The PCR conditions consisted in an initial denaturation step at 95°C for 2 min, followed by 45 cycles of denaturation at 95°C for 30 sec, annealing at 50°C for 30 sec, and extension at 72°C for 30s, followed by a final extension step at 72°C for 7 min. We included a negative PCR control and two positive controls (mock communities MC_1_ and MC_2_ including 17 arthropod taxa each; [44]). Amplicons were checked on a 2% agarose gel stained with GelRed (Biotium). Library preparation was performed by MACROGEN (Seoul, Korea) with the TruSeq Nano DNA kit and unique dual indexing (Illumina, USA). Prepared libraries were pooled at equal molarities before sequencing on two lanes of an Illumina MiSeq spiked with 30% PhiX using 150 bp paired-end chemistry.

### 2.4 Sequence data processing

We used the FROGS pipeline (“Find Rapidly OTU with Galaxy Solution”; [45]) to merge paired-end sequences and removed low-quality reads (<Q30) with VSEARCH v.2.17.0 and to trim primers with CUTADAPT v.2.10. Then, we filtered sequences by length (140-170bp) and for any ambiguities, and clustered them into operational taxonomic units (OTUs) using a maximum aggregation distance of one mutation with the SWARM algorithm [46]. Chimeras were then removed using VSEARCH [47] with de novo UCHIME [48], and a quality filter was carried out by removing any OTU occurring at a frequency below 0.005% in the whole dataset. In addition, we used isBimeraDenovo from dada2 [49,50] to remove the residual chimeric sequences which were not detected using the FROGS pipeline. Filtering for potential false positives was carried out by discarding any count of less than 5 reads.

### 2.5 Taxonomic assignment & reliability of the data

Taxonomic affiliation of OTUs was first performed using NCBI BLAST + automatic affiliation tool available in FROGS pipeline, with the Public Record Barcode Database (data related to BOLD database http://v3.boldsystems.org in February 2019, with maximum 1% of N: “COI BOLD 1percentN 22019” reference database). Multi-affiliations were manually checked and resolved with the affiliation Explorer tool on shiny migale (shiny.migale.inrae.fr). Taxonomic affiliations were then individually checked with both the NCBI BLASTn tool and BOLD database. When results were not concordant with FROGS’s output, the most probable taxa name was considered: at the species level if the sequence had ≥ 98% similarity to a single species in NCBI, at the genus level when ≥ 97%, to the family level when ≥ 92% and to the order level when ≥ 90%. When multi-affiliations occurred (*e*.*g*. different species within in a same genus), taxa name was downgraded to the common taxonomic level (*e*.*g*. genus). BOLD database was used in parallel to check for similar outputs between the two databases, and in case of unresolved multi-affiliations with NCBI, BOLD taxa names were retained when ≥ 98% similarity. To increase taxonomic determination of mosquitoes, a comparison of sequences was further done with unpublished sequences of *Culex neavei* collected in Reunion Island, because this species was absent from the reference databases. Finally, OTUs with the same affiliations were collapsed when they were present in the same samples (potentially resulting from PCR errors, heteroplasmy or intra-specific diversity). We produced two OTU tables: the first including the number of reads for each taxa in each feces, and the second detailing the presence/absence (coded as 0 and 1s) of each taxa in each feces.

We checked for appropriate sequencing depth per sample by calculating the percentage of estimated diversity covered in each sample using the function *depth*.*cov* from the R package *hilldiv* [51]. To assess the completeness of our sampling, we also created accumulation curves, at the family, genus and species levels, with 95% confidence levels based on 1000 bootstraps, using the R package iNEXT [52]. A diversity measure, based on Hill numbers, was calculated for *q* = 1 which is equivalent to Shannon diversity (which considers both richness and evenness of taxa).

### 2.6 Analysis of dietary diversity and composition

Variation in the dietary composition, at different taxonomic levels, was explored using histograms and bubble plots of the frequency of prey occurrence, computed with the *ggplot2* R package [53]. To assess the frequency of unique and specific OTUs, we calculated the percentage of OTUs detected only in one sample (i.e. “non-shared OTUs”) globally, and for each roost.

Variation of prey diversity (alpha diversity, Hill numbers: *q* = 1) was analyzed using gamma Generalized Linear Models (GLM). We first tested the effect of roost and sex (and their interaction), by including them, as well as sequencing depth, as explanatory variables. We also tested the effect of pregnancy on prey diversity, by using female data only, and including pregnancy status and roost (and their interaction), as well as sequencing depth as explanatory variables. Then, to understand the role of land-use in the distribution of prey diversity, we applied gamma Generalized Linear Mixed-Effect Models (GLMM) using *glmmTMB* package [54]. We included sex and land-use variables (as well as their interactions) as fixed effects, and roost as a random effect. However, because the semi-natural variable was highly negatively correlated to the two other land-use variables (collinearity assessed by calculating Pearson correlation coefficients), we removed it from the model and only kept urban and agricultural variables as predictors. We applied this modeling approach not only to overall prey diversity but also separately to the two dominant prey orders (Lepidoptera and Blattodea; see results). Additionally, we examined changes in the relative abundance of these prey orders using beta GLMMs, following data transformation to handle 0 and 1 values appropriately. Model specifications are detailed in Table S1. For all models, we assessed residual distributions using QQ plots generated with the *DHARMa* package and tested the significance of predictors using Type II ANOVA from the *car* package in R [55].

To assess the drivers of diet composition, a Bray–Curtis distance matrix of samples was computed based on the OTU table, using the *vegdist* function of the *vegan* package [56]. We verified homogeneity of variance among roosts and sexes using the *betadisper* function. We then performed a permutational multivariate analysis of variance (PERMANOVA) using the *adonis* function and 999 permutations, including roost and sex as predictors, and the by = “margin” argument to allow for the independent assessment of each main effect while controlling for the presence of the other in the model. To infer the role of land-use in structuring dietary composition, we also used a Mantel test to compare the diet matrix (Bray-Curtis) to the landscape dissimilarity matrix calculated above (999 permutations). To visualize the structure of dietary composition, non-metric multidimensional scaling (NMDS) ordinations were conducted on the Bray–Curtis distance matrix with the *metaMDS* function. Seven outlier samples in the initial plot (MF082, MF086, MF098, MF139, MF196, MF200 and MF298) were removed from the NMDS plots to better observe grouping patterns.

For each roost, we also calculated the percentage of OTUs that were not shared with other roosts (i.e. “roost-specific OTUs”). To identify taxa that differed significantly in relative abundance among roosts, we used a linear discriminant analysis effect size (LEfSe; Galaxy v.1.0; [57,58]). For this analysis, we set the alpha value for the Kruskal–Wallis test at 0.05 and the threshold on the logarithmic LDA score at 2.0.

## 3 RESULTS

### 3.1 Check of controls

The MiSeq sequencing run respectively produced 6 and 10 OTUs (each represented by only 1 read) for the negative controls of extraction and PCR. These contaminant reads were automatically removed by filtering steps leading to cleaned negative controls. In contrast, for the two mock communities (MCs), a total of 81588 and 110899 reads was produced after bioinformatic processing. However in MC_1_, only 6 taxa over the expected 17 were identified, keeping in mind that one expected taxa (Blattodea) was removed during filtering steps (because of only 189 reads), and that 4 taxa were already known to not be amplified by Zeale’s primers [44]. In MC_2_, 8 taxa over the expected 17 were identified (among them, 4 were already known to not be amplified by Zeale’s primers). Both positive and negative control samples were removed from the dataset for further analyses.

### 3.2 Bat data filtering and reliability

After removing controls from the dataset, a total of 8 972 457 reads from bat samples were processed. Bioinformatic processing excluded 1.2% of these reads, and led to 281 OTUs. After taxonomic assignment, 13 OTUs were discarded: 9 were absent from database (*i*.*e*., blast with FROGS produced no result) and 4 OTUs were likely to be non-specific (e.g., either nematodes or crustaceans). After chimera detection and OTU collapsing, the final dataset contained 174 OTUs in 89 bat samples (Table 1) and a mean number of 92 146 (+/-4 424, % 95CI) reads per sample. Our results revealed an appropriate sequencing depth per sample, with the estimated diversity covered in each sample always above 99%. The extrapolated accumulation curves almost reached a plateau at the family, genus and species levels (Figure S2), highlighting that our global sampling effort was probably close to capturing all prey diversity at the studied sites.

**Table 1.** Details of samples analyzed for the diet composition of Reunion free-tailed bats. *For ESA, 15 samples were initially sequenced but one (MF259) was removed after bioinformatic processing because only 2 reads were produced. In the TBA roost, huge quantities of guano are accumulating under the colony. The number of roost-enriched OTUs corresponds to the result of the Lefse analysis, comparing the abundance of OTUs in one roost to the others roosts.

| Roost name | TBA | EGI | ESA | VSP | PSR | TGI | RBL | Total |
| --- | --- | --- | --- | --- | --- | --- | --- | --- |
| Habitat | Cave | Building | Bridge | Bridge | Bridge | Building | Bridge |  |
| N fecal samples<br>(M/F) | 11<br>(0/11) | 11<br>(6/5) | 14<br>(12/2)* | 12<br>(6/6) | 13<br>(10/3) | 14<br>(8/6) | 14<br>(5/9) | **89**<br>**(47/42)** |
| Mean N reads/sample<br>(+/-95%CI) | 81173<br>(8717) | 88096<br>(14330) | 88009<br>(9274) | 92528<br>(11703) | 95217<br>(11879) | 103685<br>(18743) | 93370<br>(9361) | **92146**<br>**(4424)** |
| N OTUs | 14 | 35 | 55 | 54 | 64 | 32 | 47 | **174** |
| N roost-specific OTUs<br>(%) | 6<br>(43%) | 21<br>(60%) | 12<br>(22%) | 14<br>(26%) | 35<br>(55%) | 7<br>(22%) | 10<br>(21%) | **105**<br>**(60%)** |
| N roost-enriched OTUs<br>(%) | 7<br>(50%) | 21<br>(60%) | 19<br>(35%) | 18<br>(33%) | 42<br>(66%) | 8<br>(25%) | 17<br>(36%) | **132**<br>**(76%)** |
| N non-shared OTUs<br>among samples<br>(%) | 13<br>(93%) | 30<br>(86%) | 38<br>(69%) | 36<br>(67%) | 37<br>(58%) | 21<br>(66%) | 27<br>(57%) | **83**<br>**(48%)** |

### 3.3 Taxonomic identity of prey

From the 174 OTUs obtained, 169 (97%) were taxonomically identified at the order level (5 OTUs were classified as unknown Insecta), 102 (59%) at the family level, 64 (37%) at the genus level, and only 37 of them (21%) at the species level. Globally, most prey belonged to Lepidoptera (48% of reads, 45% of OTUs; Figure 2), which were highly diverse as we detected the largest proportion of sequence diversity in this group (75 different OTUs). Lepidopterans were largely consumed in all roosts, except in TBA (< 1% of the reads) where Hemiptera, instead, were the predominant prey. Blattodea and Diptera were also globally abundant in the dietary composition of bats, encompassing 17% and 15% of reads (10% and 20% of OTUs), respectively (Figure 2).

**Figure 2.**
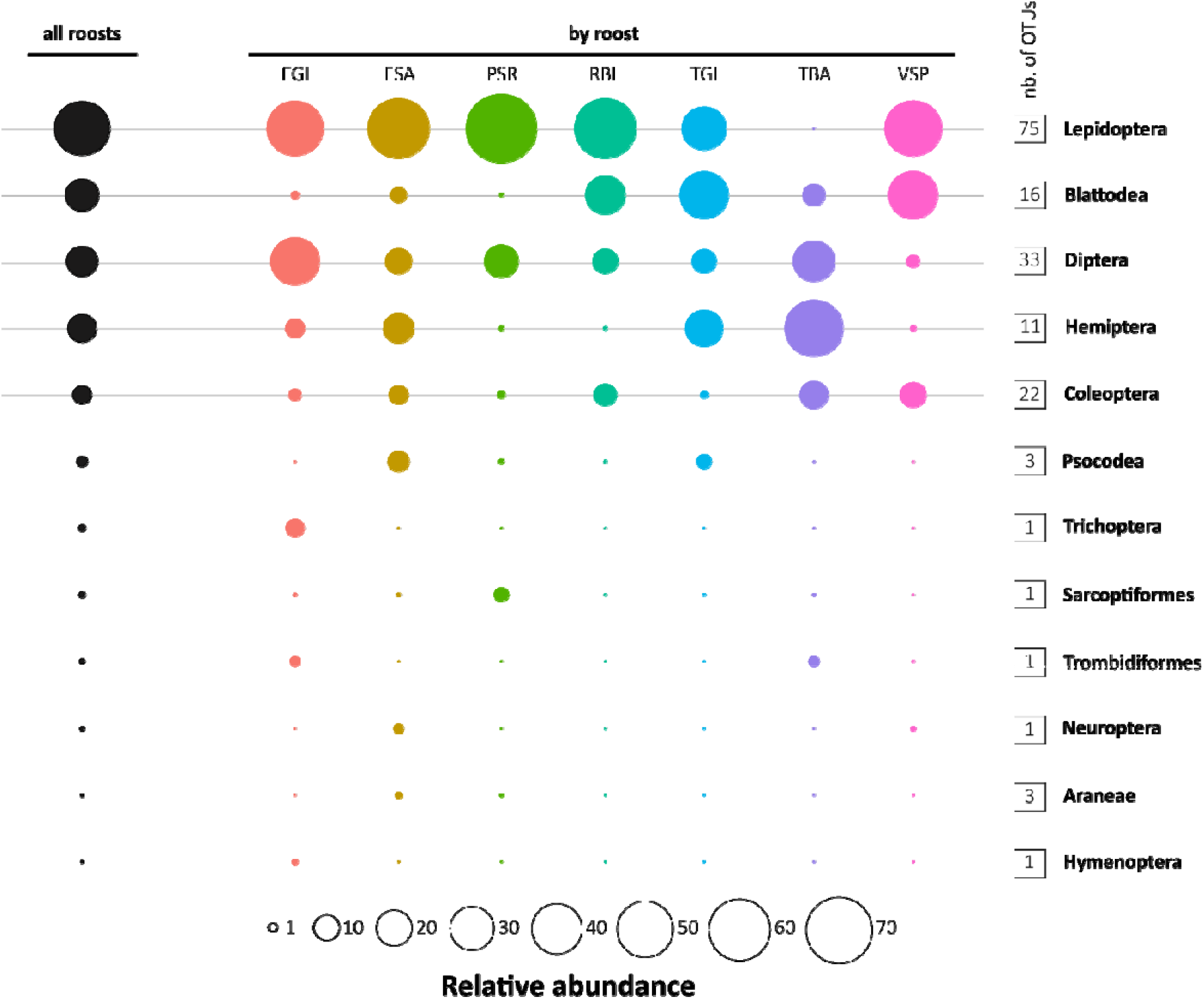
Relative abundance of the 12 arthropod orders taxonomically identified in the diet of Reunion free-tailed bats. Relative abundance was calculated over the whole dataset (left) and within each roost (right), excluding 6 OTUs that were unclassified at the order level. The number of OTUs in each arthropod order is indicated in squares on the right.

Among the 174 OTUs, only 12 were commonly detected (*i*.*e*. at least present in 10% of samples) and most of them belonged to Lepidoptera (Figure 3). The most frequent species were an unclassified Lepidoptera (present in 35% of samples) and an unclassified Chrysopeleiinae (in 33% of samples; Figure 3). Among Blattodea, an unclassified Kalotermidae1 and *Blattella lituricolis* were the two taxa the most frequently observed (Figure 3).

**Figure 3.**
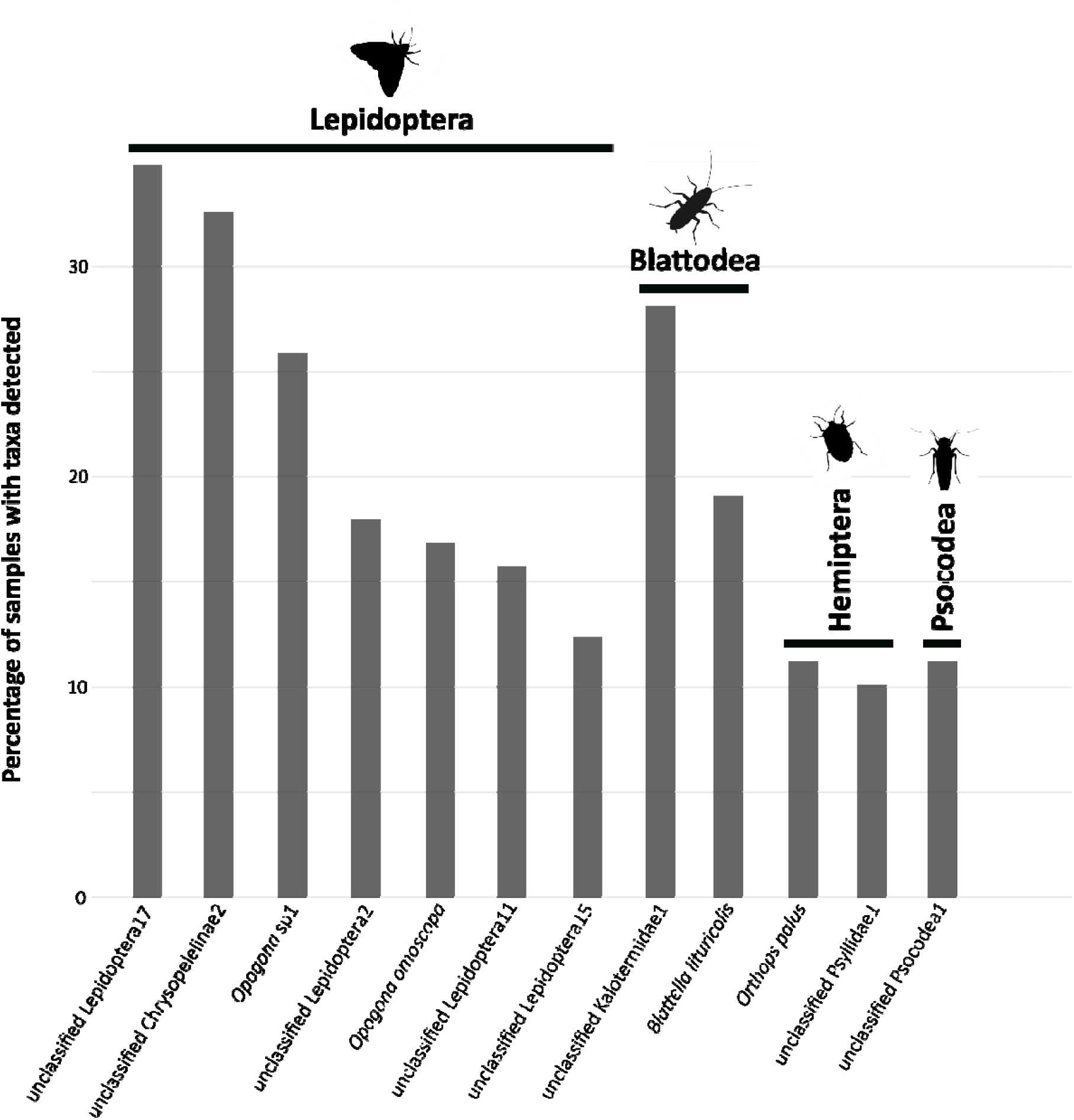
Commonly detected taxa species in the diet of Reunion free-tailed bats. Species shown are OTUs present in at least 10% of samples and organized by arthropod order.

We detected several taxa of interest in the diet of Reunion free-tailed bats, including vegetal pests and animal disease vectors (Table 2). For example, the bug *Orthops palus* is considered as a major pest of mango in Reunion Island, and it also has a broad range of plant hosts, such as avocado and lychee [59]. We detected this insect in the feces of ten bat individuals from four different roosts. We also identified two species of invasive termites (*Cryptotermes brevis* and *Cryptotermes havilandi*) which are serious pests that cause damage to wooden structures [60]. Although they were present in low abundance in the diet of Reunion free-tailed bats, four mosquito species were also identified, including *Aedes albopictus, Aedes fowleri, Culex quinquefasciatus* and *Culex neavei*. In addition, a hematophagous midge of the genus *Culicoides* was identified, although present in a limited number of bat samples (n = 4). Interestingly, Reunion free-tailed bats also fed on non-flying arthropods such as spiders, including two distinct species of *Cheracanthium*.

**Table 2.** Examples of taxa of interest present in the diet of Reunion free-tailed bats.

| Species | Nb. bat samples | NCBI ID percentage |
| --- | --- | --- |
| <u>Crop pests</u> |  |  |
| *Opogona sacchari* | 5 | 100 |
| *Orthops palus* | 10 | 100 |
| *Sternochetus mangiferae* | 1 | 100 |
| <u>Wood termites</u> |  |  |
| *Cryptotermes brevi* | 4 | 100 |
| *Cryptotermes havilandi* | 1 | 100 |
| <u>Mosquitoes</u> |  |  |
| *Aedes albopictus* | 2 | 100 |
| *Aedes fowleri* | 1 | 100 |
| *Culex quinquefasciatus* | 2 | 100 |
| *Culex neavei* | 4 | 100 |
| <u>Biting midges</u> |  |  |
| *Culicoides* sp. | 4 | 98.7 |
| <u>Spiders</u> |  |  |
| *Cheirachantium insulare* | 2 | 100 |
| *Cheirachantium insigne* | 1 | 100 |

### 3.4 Drivers of dietary diversity and composition

On average, we recovered 5.9 ± 0.9 prey species per fecal sample (min = 1, max = 20). We found that the species-level dietary diversity (Hill number: *q* = 1) was significantly different among roosts (GLM_1_: χ^2^_6_ = 33.148, *P* = 9.819^-06^; Figure 4a). Globally, we found weak evidence that females consumed a larger diversity of prey than males (GLM_1_: χ^2^_1_ = 3.875, *P* = 0.049; Figure S4a), and no effect of pregnancy on the diversity of prey consumed by females (GLM_2_: χ^2^_1_ = 1.285, *P* = 0.257; Figure S5). However, at a lower taxonomic level, we found that males consumed more Lepidoptera (GLMM_4_: χ^2^_1_ = 7.041, *P* = 0.008; Figure S4b), but less diverse Blattodea (GLMM_7_: χ^2^_1_ = 13.125, *P* = 0.0003; Figure S4c) than females.

**Figure 4.**
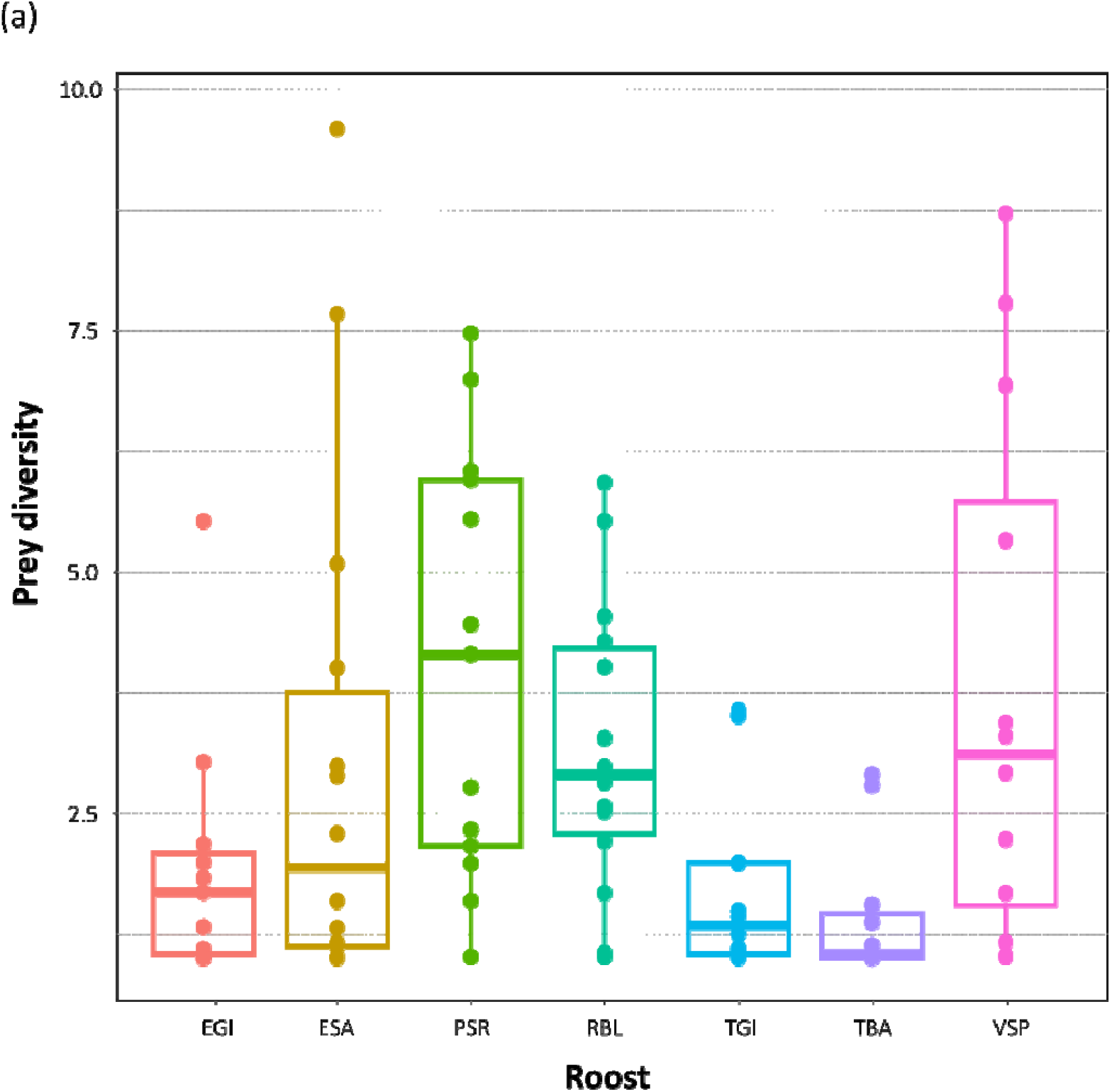
Spatial variation of the diet of Reunion free-tailed bats. Observed arthropod diversity in the diet of bats from seven roosts. Diversity was calculated based on Hill numbers and *q* = 1.

PERMANOVA analyses revealed significant variation of species-level dietary composition among roosts (*P* = 0.001) but not sex (*P* = 0.219). These results were supported by the LEfse analysis which identified 132 OTUs (76% of total OTUs) that were differentially distributed among roosts (Figure S3). Dietary variation among roosts was also illustrated by the large proportion (60%) of roost-specific OTUs (Table 1). However, NMDS plots showed some overlap among roosts (Figures 5 and S6) and highlighted the variation among bat individuals, as well as the presence of rare prey (outliers) which probably exacerbated the dissimilarity between roosts. Indeed, this variation among bat individuals was supported by the fact that 48% (n = 83) of OTUs were sample-specific (*i*.*e*. detected in only one sample), and those were particularly abundant in the TBA and EGI roosts (93 and 86% of sample-specific OTUs in these roosts).

**Figure 5.**
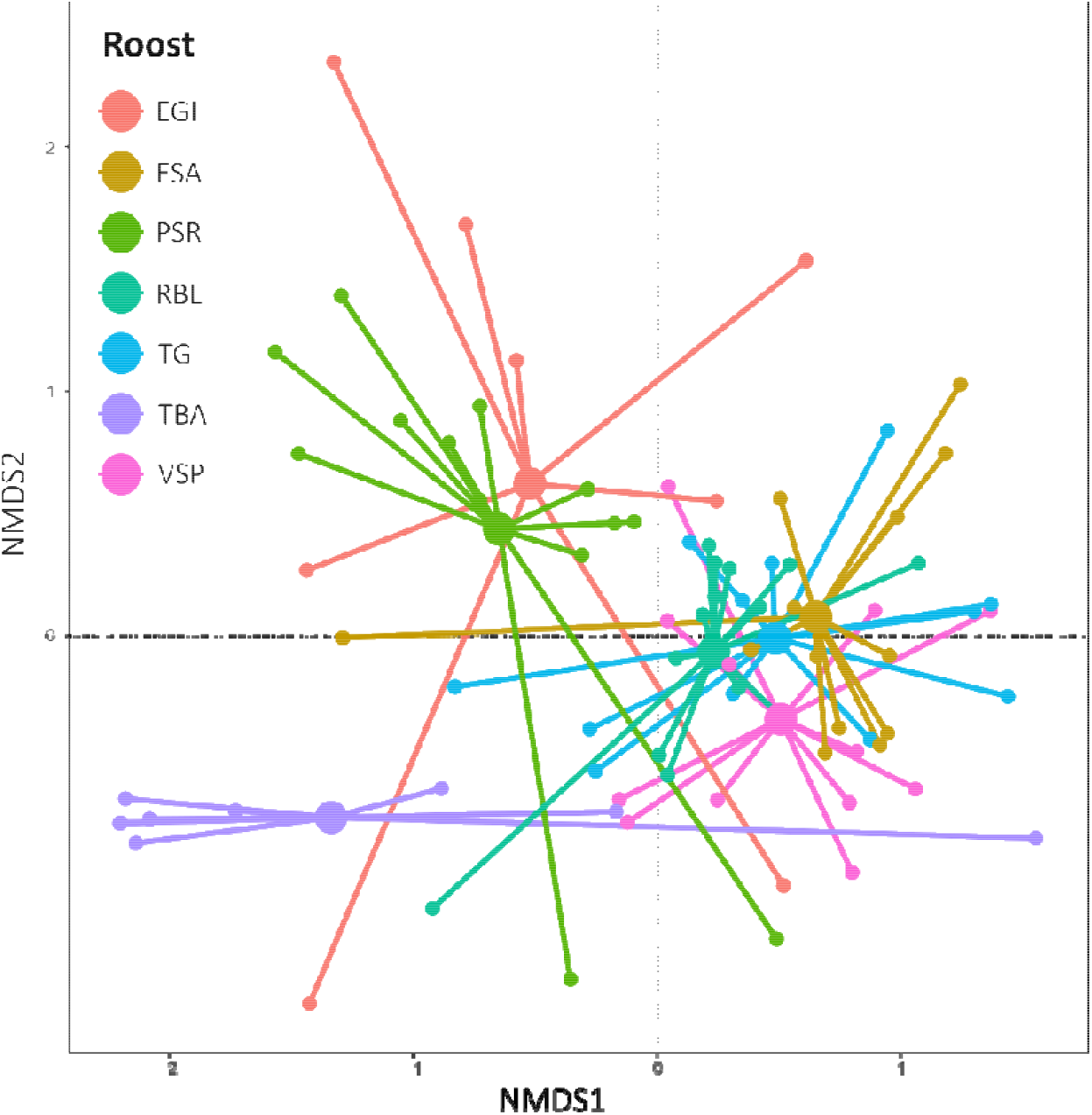
Non-metric multidimensional scaling (NMDS) ordination plot of the diet composition of Reunion free-tailed bats, according to the sampling roost. Large circles represent group centroids, while small circles correspond to individual fecal samples, connected to their respective centroids by lines. Plot was produced after removing seven outlier samples for better visualization. NMDS plot including the whole dataset is available in Figure S4.

The PCA showed that our sampled roosts had a different landscape composition, based on a matrix of three land-use variables (urban, semi-natural and agriculture; Figure 1b). We found evidence that landscape surrounding roosts was correlated to the dietary composition of bats (Mantel test: r^2^ = 0.197; *P* = 0.001; Figure S7), but not to the global prey diversity that the bats consumed (GLMM_3-agri_: χ^2^_1_ = 2.448, *P* = 0.118, GLMM_3-urban_: χ^2^_1_ = 0.023, *P* = 0.881). At a lower taxonomic level, agriculture was associated with the consumption of more Lepidoptera (GLMM_4_: χ^2^_1_ = 5.330, *P* = 0.021; Figure 6a), and slightly more diverse Blattodea (GLMM_7_: χ^2^_1_ = 3.976, *P* = 0.046; Figure 6b). In urban areas, bats consumed significantly more Blattodea (GLMM_6_: χ^2^_1_ = 8.698, *P* = 0.003; Figure 6c). This is illustrated by the high abundance of wood termites and cockroaches in the diet of bats roosting in highly urbanized areas, such as in TGI, VSP and RBL roosts (Figure 6d).

**Figure 6.**
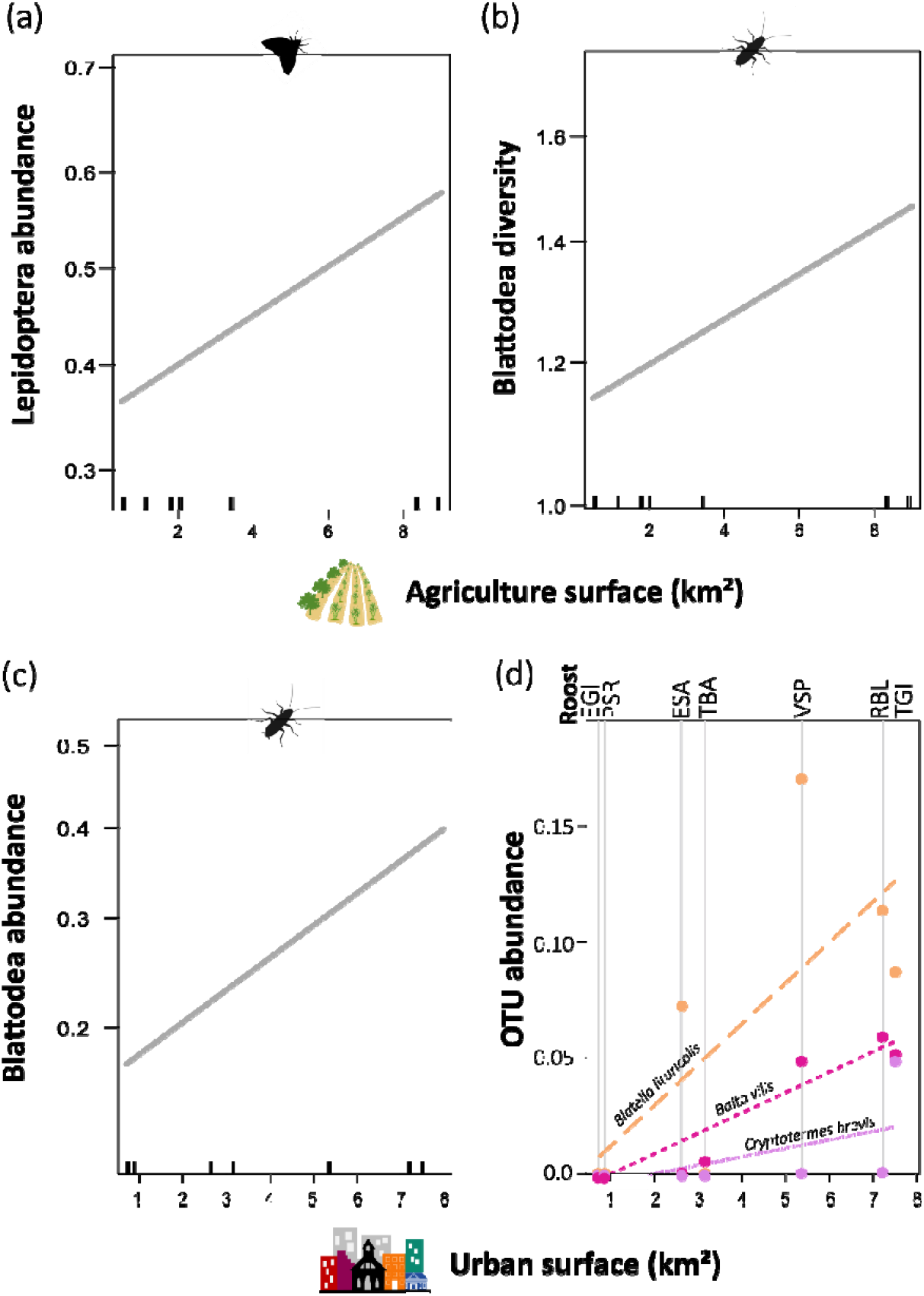
Variation of prey abundance (a,c,d) and diversity (b) in Reunion free-tailed bats roosting in agricultural and urban landscapes. Diversity was calculated based on Hill numbers and *q* = 1. For (a-c), predictions are based on models in Table S1 and shaded areas depict standard errors below and above the estimated mean responses. For (d), OTU abundance was calculated from raw data per roost, for a wood termite and two cockroach species (Blattodea). To enhance visualization, circles corresponding to identical zero values were slightly jittered.

## 4 DISCUSSION

In this study, we provided a first description of the Reunion free-tailed bat diet. Our quantification of dietary diversity (174 OTUs from 12 arthropod orders) was probably underestimated as (i) extrapolated accumulation curves suggested that a greater sampling effort could reveal more prey at the genus and species levels, and (ii) certain taxa were not retrieved from the mock communities, suggesting failure in PCR amplification. Our study was also based on the examination of feces samples collected during a small-time frame (39 days), thus representing only a small fraction of the biological cycle of this bat species. Because of differences in bats’ energy requirements through time and seasonal changes in prey availability [24], samples collected during other times of the year are needed to capture the entire prey community [61]. Finally, although resource partitioning likely facilitates the coexistence of bat species in highly diverse communities [62,63], only two insectivorous bat species are present in Reunion Island [64]. The second one, *Taphozous mauritianus*, is far less abundant than free-tailed bats, probably lowering interspecific competition for insects, and thus favoring a wide dietary niche for Reunion free-tailed bats.

Our results revealed that Reunion free-tailed bats fed on several agricultural and household pests, and at a lesser extent on animal disease vectors. Agricultural pests include, but are not limited to, those affecting mango, avocado and lychee, which are economically important sectors in Reunion Island. Because we were limited in identifying prey to the species level (only 21% of OTUs), further analyses, based on a broader and more accurate reference database, are needed to resolve species identification. This will probably lead to the identification of additional agricultural pests, especially among the Lepidopterans which are predominant in our study. Household pests include two species of termites. Considering animal disease vectors, Reunion free-tailed bats consumed the tiger mosquito *Aedes albopictus*, the dengue and chikungunya vector in Reunion Island [65], although it was not a major prey in our study. This is consistent with other studies where mosquitoes represent only a small proportion of bat diet [20] and is explained by the biology of this mosquito species, which is more diurnal [66]. Three other mosquito species, which are known to bite humans, have been identified, including *C. quinquefasciatus* which is a night-active mosquito very aggressive towards humans, and *C. neavei* and *A. fowleri*. Although not associated to human diseases in Reunion Island, these three species are known to transmit arboviruses such as Usutu, Rift Valley Fever and West Nile viruses in other parts of the world, including in the neighboring islands of the Western Indian Ocean (Madagascar, Comoro archipelago, etc.) [67–69]. One species of biting midge, which remains to be classified at the species level (*Culicoides* sp.), was also identified and is of importance as this genus is known to transmit bluetongue and epizootic hemorrhagic disease viruses to ruminants [70].

Because of the wide distribution of Reunion free-tailed bats over the island and their large population size [39,40], our results suggest that this species could play a significant role in reducing insect crop damage and household pests. Additionally, it may help mitigate vector-borne diseases and nuisance insects that affect both humans and animals. Similar benefits have already been demonstrated in various insectivorous bat species [18,20,71–74]. For example in Italy, the presence of bats in apple orchards reduces by 50% the weight of apples per tree damaged by the codling moth *Cydia pomonella* [75]. Enclosure experiments have also demonstrated that the northern long-eared bats (*Myotis septentrionalis*) may prey on ovipositing *Culex* mosquitoes, contributing to a 32% reduction in egg-laying [76]. Protecting Reunion free-tailed bat colonies, together with erecting artificial bat houses to attract bats, may therefore reduce the use of pesticides and provide economic benefits [72,77]. Reunion free-tailed bats thus represent a natural asset in agroecology and epidemiology, for regulating economically important insects, as well as for public and animal health issues.

Reunion Island’s landscape has changed dramatically due to recent human colonization, which may profoundly affect insect communities and shape insectivorous bat’s diet. Indeed, our findings revealed that the diet of Reunion free-tailed bats was closely linked to these landscape changes, and that agriculture and urbanization were associated with differences in dietary diversity and composition. Contrary to our predictions, we did not evidence a global decrease of diet diversity in agricultural landscapes, but showed in contrast an increase of abundance and diversity for specific prey orders (Lepidoptera and Blattodea, respectively). Our results differed from those found for the common bent-wing bat (*Miniopterus schreibersii*), which reduces its dietary diversity in intensive agriculture areas [26], and urban noctule bats (*Nyctalus noctula*) which displayed a more diverse diet than their rural counterparts [71]. In Reunion Island, agriculture is dominated by sugar cane [78] and for our study sites, sugar cane represents up to 65% of the agriculture areas surrounding bat roosts. Sugar cane fields are often associated with many moth species [79], which is consistent with the increased consumption of Lepidoptera for bats roosting in agricultural landscapes. As previously shown for *M. schreibersii*, which specializes on the moths present in the local crop systems across Southern Europe [26], our study suggested an opportunistic behavior of Reunion free-tailed bats to feed upon moths in sugar cane fields. This is in line with the selection of sugar cane habitats for foraging in two African molossid bats (*Chaerephon pumilus* and *Mops condylurus*) [80]. Exclusion experiments could be designed to assess whether Reunion free-tailed bats exert sufficient predation pressure to offer ecological and economic benefits to agroecosystems. Indeed, such experimental studies have shown a significant reduction in crop damage due to the presence of insectivorous bats (e.g. [19,81–83]), and would thus help evaluating the potential role of Reunion free-tailed bats in sustainable pest control approaches.

Despite the general negative effect of urbanization on insect communities [84], we did not observe significant changes in overall dietary diversity among bats roosting in urban areas. The maintenance of dietary diversity in urban-roosting bats could result from the high mobility of this bat species [39,40], allowing exploitation of resources beyond urban centers. Indeed, urbanization selects highly mobile species with more generalist diet that are better able to exploit available resources [85]. A similar pattern has been recently suggested in urban common noctule bats, which commute to the outskirts of cities to forage over farmland and forested areas [86]. By doing so, they not only prey on urban insect species but also on rural species. This foraging flexibility may compensate for the general decline in urban insect diversity and, in the case of urban common noctule bats, even lead to a more diverse diet among bats roosting in urban areas [71]. Our findings also indicate that urbanization could drive predation of Blattodea, including several species of wood termites and cockroaches. This shift likely results from the increased availability of these synanthropic insects, which thrive in human-modified environments due to abundant food waste, decaying wood, and artificial shelters, making them easily accessible prey. Together, these findings highlight the dietary plasticity of Reunion free-tailed bats in human-modified landscapes. To further elucidate their feeding ecology, future research should compare their diet with insect availability across different landscapes on Reunion Island. This approach will help confirm whether landscape is driving dietary specialization or if bats maintain a generalist foraging strategy and hunt opportunistically.

Although intraspecific variations in bat diet composition are rarely considered, sex- and reproduction-related differences have been documented in several bat species, including molossids like the European free-tailed bat *Tadarida teniotis* [28]. In Reunion free-tailed bats, longitudinal sampling showed that sexual segregation for roosting is common, especially during gestation periods, suggesting different ecological requirements between sexes [39]. Our findings indicate that males globally consume more Lepidoptera than females, and a higher diversity of Blattodea. These differences may stem from varying prey preferences in response to different energy requirements and intraspecific competition [27– 29]. In our study, bats were sampled during the gestation period but we could not detect any effect of pregnancy on diet diversity. This contrasts with previous findings on female pond bats [29], which may partly be explained by differences in the reproductive stages represented in the samples. Notably, the earlier study included both pregnant and lactating females, whereas our dataset did not include lactating individuals, whose energetic demands are typically higher and may drive more pronounced dietary shifts. Additionally, a limited sample size and potential overlap in foraging habitats between reproductive and non-reproductive individuals due to communal roosting, may limit dietary flexibility. Therefore, future radio-telemetry studies are crucial to track the movements of Reunion free-tailed bats. This will help confirm whether, despite roosting together, females and males of Reunion free-tailed bats use distinct habitats to forage [87], and whether sex-specific dietary strategies and responses to urbanization occur in this bat species.

Considered as common city-dwellers, Reunion free-tailed bats provide an original model to show how some bats, thought to be well-adapted to urban life, can shift their realized dietary niche. However, dietary plasticity in bats may bring new selective pressures and thus lead to important trade-offs for bat evolution [11]. For example, a previous study suggested that *Pipistrellus kuhlii* has undergone an increase in skull size in response to urbanization, which would be adaptive to handle larger prey (moths) present in artificially illuminated cities [88]. Thus, understanding the consequences of dietary plasticity and shifts on bat health and fitness will be necessary for urban evolutionary research.

## AUTHOR CONTRIBUTIONS

M.D., B.R., H.D. and C.L. participated in the conception and design of this work. M.D., G.LM., G.D., N.H-H and C.L. were responsible for sample collection in the field. M.D., M.T. and M.G. performed laboratory work. M.D. analyzed the data, with the help of D.A.W. and M.G. M.D. supervised the project and wrote the first draft of the manuscript. All authors revised the manuscript.

## DATA AVAIBILITY STATEMENT

The list of samples, the Illumina COI reads and the OTU table have been deposited in Zenodo (10.5281/zenodo.15349409).

## ACKNOWLEDGEMENTS

We are grateful to Eco-Med Océan Indien, Biotope, the DEER of Région Réunion (Direction de l’Exploitation et de l’Entretien des Routes), the DRT of Département Réunion (Direction des Routes et des Transports of the Department Reunion), and the City hall of Salazie for their help in identifying and accessing bat roosts. We also thank Axel Hoarau, Céline Toty and Pablo Tortosa for their assistance in the field and Brice Derepas for help with spider identification. We also thank Samuel Nibouche for his help with taxon identification and input for statistical analyses. We are grateful to the genotoul bioinformatics platform Toulouse Midi-Pyrénées, and Sigenae group for providing help and computing resources thanks to Galaxy instance (http://sigenae-workbench.toulouse.inra.fr). This research was supported by the French National Research Agency (ANR JCJC SEXIBAT) and by the Université de la Réunion BioST (MOLOSS-EAT).

## ETHICS STATEMENT

Bat capture and manipulation were evaluated by the animal ethics committee of Reunion Island, approved by the French Ministry of Research (APAFIS#10140-2017030119531267), and conducted under a permit delivered by the Direction de l’Environnement, de l’Aménagement et du Logement (DEAL) of Reunion Island (DEAL/SEB/UBIO/2018-09).

