## Supplementary Information for "Urban bats change the menu: dietary plasticity across human-modified landscapes of a tropical island"

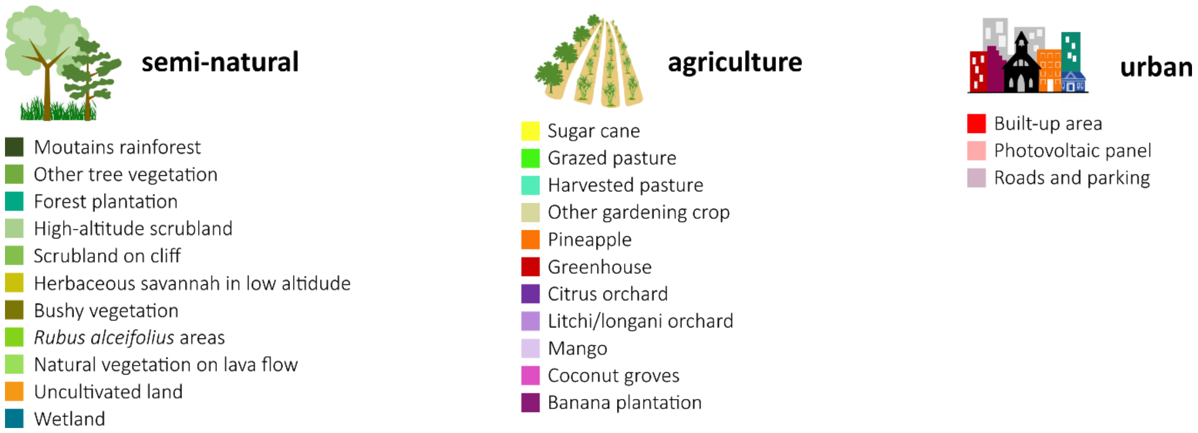


**Figure S1. List of habitats included in the three land-use variables.** Colors refer to those in the map in Figure 1.

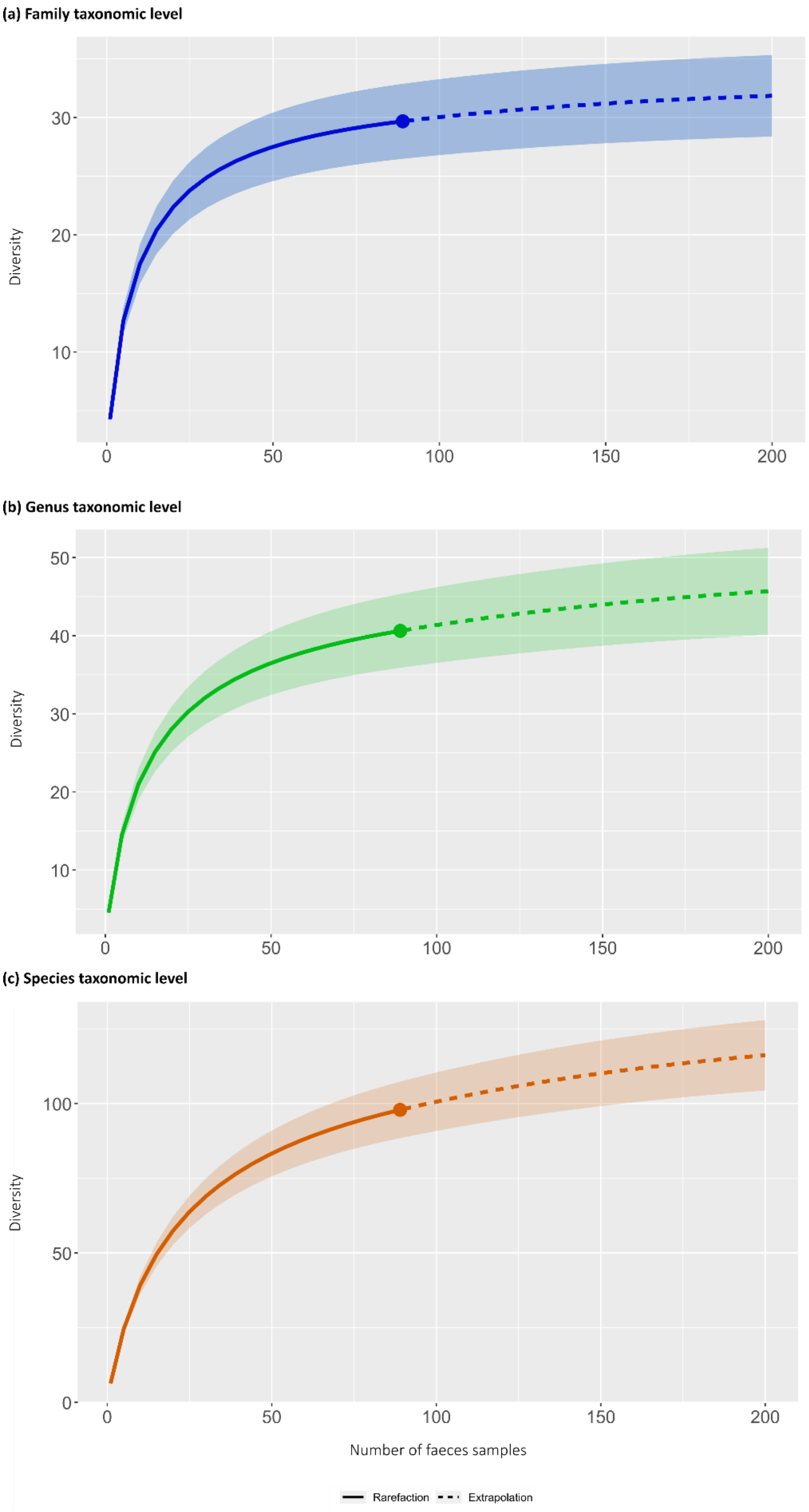


**Figure S2. Accumulation curves (and 95% confidence intervals) of prey diversity estimated by Hill numbers (q=1, equivalent to the Shannon index estimator) at the (a) family, (b) genus and (c) species levels.**

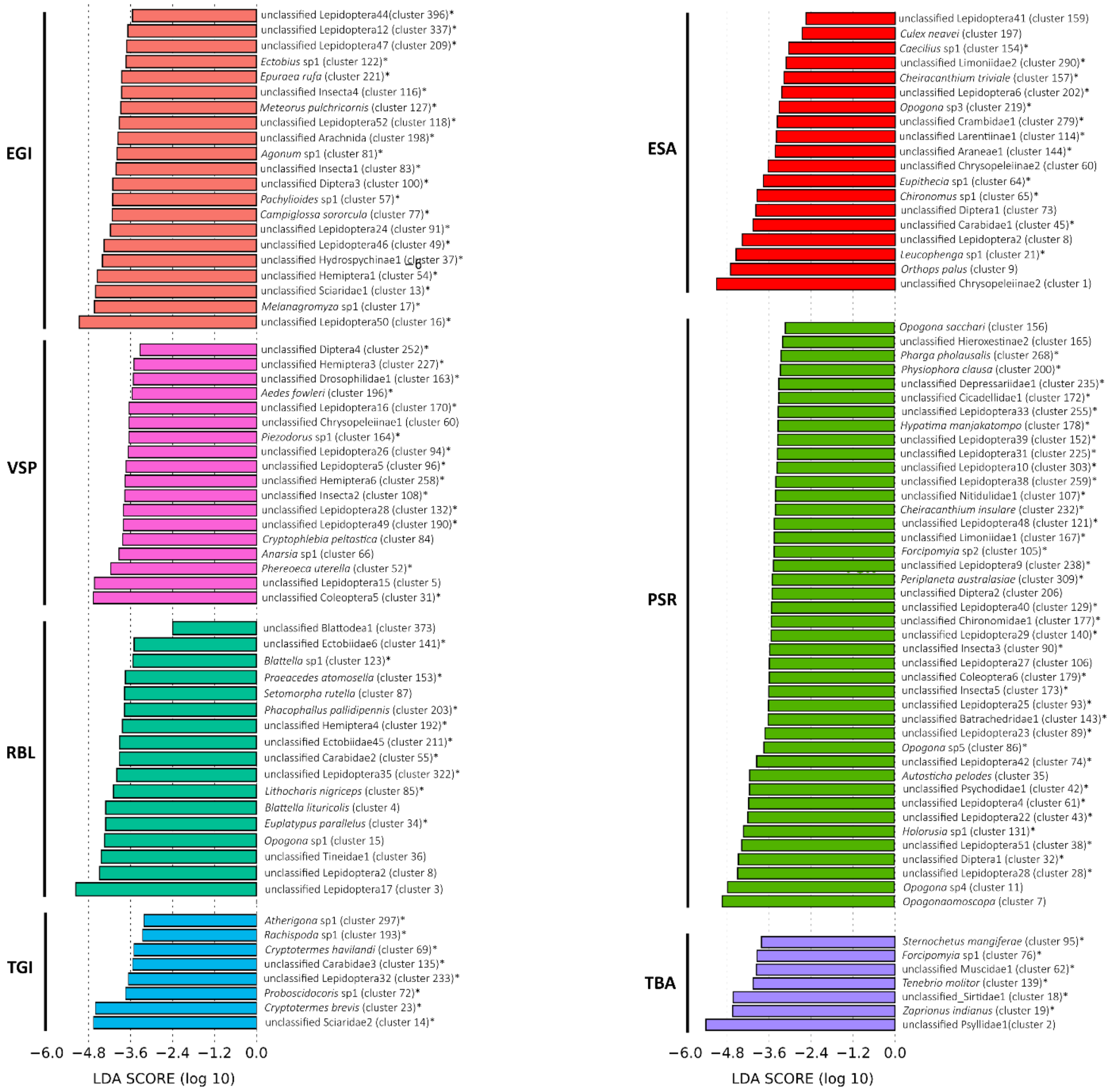


**Figure S3. LEfse analysis showing OTUs significantly more consumed in seven roosts of Reunion free-tailed bats.** OTUs with an asterisk were only detected in the specific roost.

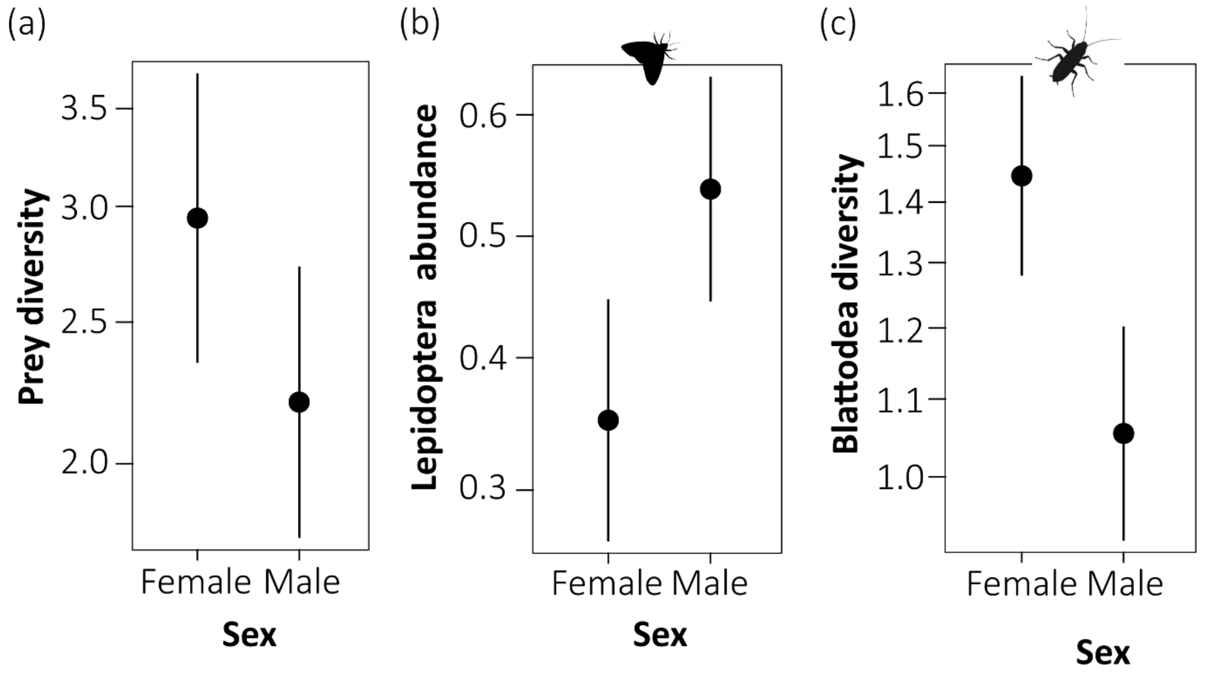


**Figure S4. Variation of prey diversity and abundance in Reunion free-tailed bats according to sex.** Predictions are based on models in Table S1. Bars depict standard errors below and above the estimated mean responses.

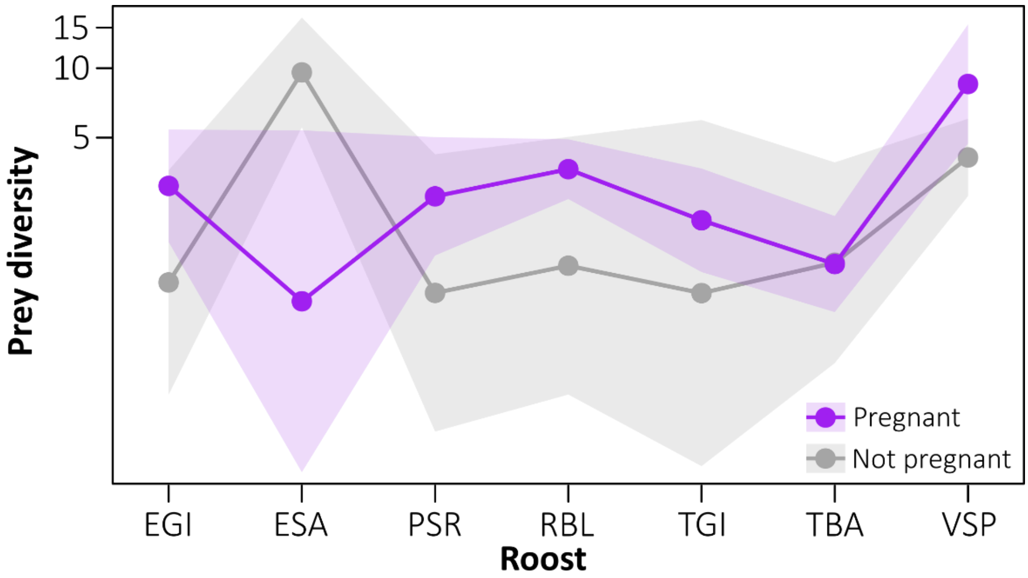


**Figure S5. Variation of prey diversity in Reunion free-tailed bats according to pregnancy status.** Raw observed data are presented in the different sampled roosts. Shaded areas depict standard errors below and above the estimated mean responses.

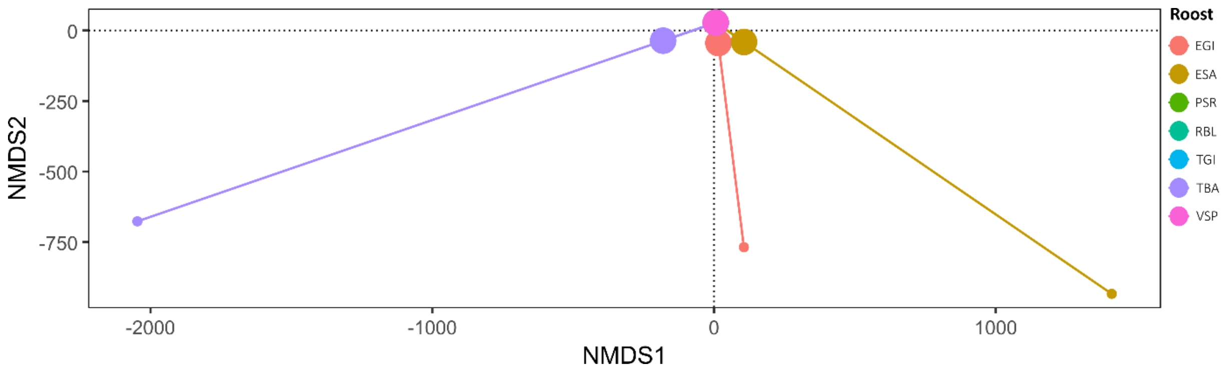


**Figure S6. Non-metric multidimensional scaling (NMDS) ordination plot of the diet composition of Reunion free-tailed bats according to the sampling roost.** Large circles represent group centroids, while small circles correspond to individual fecal samples, connected to their respective centroids by lines. Some centroids and samples may be obscured due to overlap.

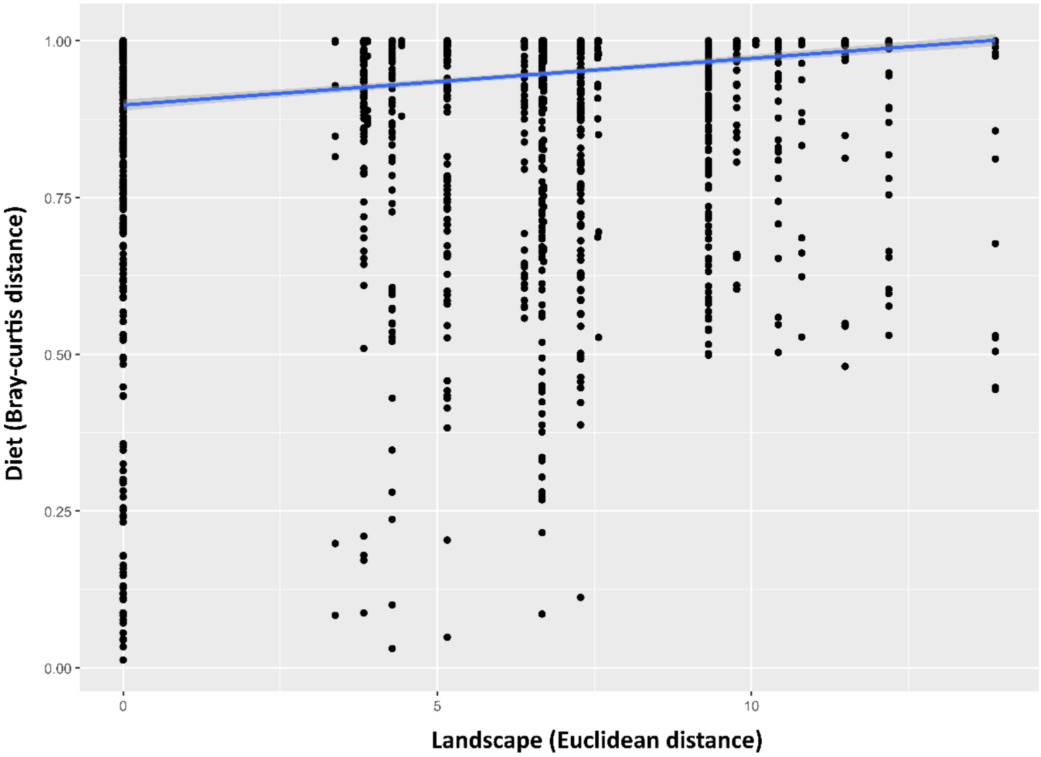


**Figure S7. Correlation of the diet (Bray-Curtis distance) and the landscape (Euclidean distance) dissimilarity matrices.**

**Table S1. Details of the GLMs used to analyse the diet of Reunion free-tailed bats.** Prey diversity was calculated using alpha diversity, Hill numbers: q=1 For prey abundance, data were adjusted with the following transformation: value * (n - 1) + 0.5) / nb.samples. For GLMM6, the model residuals showed a significant deviation (KS test: $P = 1^{e-05}$). The interaction between two variables is represented with a star and random factor in parentheses. Significant variables are underlined and highlighted in bold. Variables names are coded as follow: "Depth" = sequencing depth, "Urban" = urban surface, "Agri" = agricultural surface.

| *Model number* | *Dataset* | *Response variable* | *Explanatory variables* | *Df* | *LR Chisq* | *P* |
|---|---|---|---|---|---|---|
| $GLM_1$ | All individuals<br>N = 89 | Prey diversity | Depth<br>**<u>Roost</u>**<br>**<u>Sex</u>**<br>Roost*Sex | 1<br>**<u>6</u>**<br>**<u>1</u>**<br>5 | 0.621<br>**<u>33.148</u>**<br>**<u>3.875</u>**<br>4.483 | 0.430<br>**<u>$9.819^{-06}$</u>**<br>**<u>0.049</u>**<br>0.482 |
| $GLM_2$ | Females<br>N = 42 | Prey diversity | Depth<br>**<u>Roost</u>**<br>Pregnancy | 1<br>**<u>6</u>**<br>1 | 0.093<br>**<u>28.056</u>**<br>1.285 | 0.761<br>**<u>$9.171^{-05}$</u>**<br>0.257 |
| $GLMM_3$ | All individuals<br>N = 89 | Prey diversity | Sex<br>Urban<br>Agri<br>Sex*Urban<br>Sex*Agri<br>(Roost) | 1<br>1<br>1<br>1<br>1 | 2.998<br>0.023<br>2.448<br>1.341<br>0.900 | 0.083<br>0.881<br>0.118<br>0.247<br>0.343 |
| $GLMM_4$ | All individuals<br>N = 89 | Lepidoptera abundance | **<u>Sex</u>**<br>Urban<br>**<u>Agri</u>**<br>Sex*Urban<br>Sex*Agri<br>(Roost) | **<u>1</u>**<br>1<br>**<u>1</u>**<br>1<br>1 | **<u>7.041</u>**<br>0.307<br>**<u>5.330</u>**<br>2.815<br>0.260 | **<u>0.008</u>**<br>0.580<br>**<u>0.021</u>**<br>0.093<br>0.610 |
| $GLMM_5$ | All individuals<br>N = 89 | Lepidoptera diversity | Sex<br>Urban<br>Agri<br>Sex*Urban<br>Sex*Agri<br>(Roost) | 1<br>1<br>1<br>1<br>1 | 2.844<br>0.711<br>1.324<br>0.283<br>0.537 | 0.097<br>0.399<br>0.250<br>0.595<br>0.464 |
| $GLMM_6$ | All individuals<br>N = 89 | Blattodea abundance | Sex<br>**<u>Urban</u>**<br>Agri<br>Sex*Urban<br>Sex*Agri<br>(Roost) | 1<br>**<u>1</u>**<br>1<br>1<br>1 | 0.420<br>**<u>8.698</u>**<br>0.934<br>1.271<br>0.151 | 0.517<br>**<u>0.003</u>**<br>0.334<br>0.260<br>0.698 |
| $GLMM_7$ | All individuals<br>N = 89 | Blattodea diversity | **<u>Sex</u>**<br>Urban<br>**<u>Agri</u>**<br>Sex*Urban<br>Sex*Agri<br>(Roost) | **<u>1</u>**<br>1<br>**<u>1</u>**<br>1<br>1 | **<u>13.125</u>**<br>2.395<br>**<u>3.976</u>**<br>0.451<br>0.412 | **<u>0.0003</u>**<br>0.122<br>**<u>0.046</u>**<br>0.502<br>0.521 |
